# A quantitative approach to species occupancy across communities: the co-occurrence-occupancy curve

**DOI:** 10.64898/2026.03.19.712854

**Authors:** Vicente J. Ontiveros, Simone Mariani, Alberto Megías, Leonardo Aguirre, José A. Capitán, David Alonso

**Author notes:** **Corresponding author: David Alonso**.

## Abstract

Species tolerating the same environmental conditions can potentially colonize and thrive in the same habitats and eco-regions. Are any pair of those species equally probable to co-occur in the same community? Can we quantify the propensity of two species to co-occur together? Here, we focus on a simple but largely overlooked community-level pattern: the co-occurrence–occupancy curve, which relates the tendency of species to co-occur with others to their total occupancy across sites. We first define this empirical curve and then derive its expected shape under a random null model that assumes site equivalence and species independence. Building on these results, we introduce the Species Association Index (SAI), an occupancy-standardized measure that quantifies the tendency of a species to associate with others independently of its overall frequency of occurrence. The SAI enables meaningful comparisons among species with contrasting occupancies and provides a transparent benchmark against which departures from neutrality can be assessed. We illustrate the approach using two contrasting systems—tropical rain forest trees on Barro Colorado Island and organisms from Mediterranean rocky shores—highlighting both the generality of the co-occurrence–occupancy framework and its limitations.

## Introduction

The vast majority of research in community ecology rely on a specific mode of scientific inference, in which expectations about community structure are generated using a top-down perspective [20]. It is widely recognized that, as more information is incorporated into mechanistic and statistical models, their predictions generally become more accurate. In theory, this strategy should provide the most reliable insights into where and when species are present or absent—the lower-level pattern of interest. However, it typically requires extensive data on traits at both individual and species levels, along with information on their interactions, functional roles, dispersal pathways, habitat preferences, and related features (the upper-level complexities of individuals, species, and their interactions). In an era of rapidly advancing AI-driven statistical methods, this top-down approach can potentially result in high predictive performance, yet it offers only a limited understanding of the causal mechanisms and processes that generate the observed patterns. In this contribution, we instead use a bottom-up inferring approach. Here we first define the co-occurrence-occupancy curve—a community-level pattern to our knowledge overlooked—and then derive its expected shape under the lower-level, simplest assumptions.

Organisms from different species share the same environment in all ecological communities, including some of the harshest areas on the planet, where their tendency to co-occur may even increase as conditions become more severe [23]. The joint presence of two species in the same place and time during their life cycles can be shaped by a wide range of factors, from purely stochastic processes to extremely tight biological associations (e.g., coral–zooxanthellae symbioses). The analysis of co-occurrence data has a long tradition in ecology [35], spanning from the early, simple indices developed by Pielou and Pielou [28], through Diamond’s influential contribution [10]—and the subsequent controversies it generated [8, 11]—to more recent approaches that rely on null models to infer community assembly mechanisms [13, 14, 34]. Despite this extensive body of research, we believe that species co-occurrence is still an underexploited source of insight in community ecology. [5].

Null models are commonly used to detect cases in which species co-occur either more or less frequently than expected under random assembly [13]. Stochasticity indeed plays a major role in determining whether individuals of a species occupy a particular site, especially when habitat filtering imposes only weak constraints on genotypes (*sensu* Keddy [20]), or when community structure is largely unsettled or strongly immigration-driven, such as immediately after disturbance [24, 16, 1, 2]. Nonetheless, across ecosystems worldwide, habitat filtering, interspecific interactions, and other species-specific traits can generate systematic departures from neutrality. Quantifying these natural departures is essential for meaningful comparisons across species and communities, and for gaining insight into the mechanisms that drive community assembly. As a starting point, we must ask: what species-co-occurrence patterns should we expect under a fully random null model?

In broad terms, two main factors shape patterns of species co-occurrence: first, how common or rare each species is; and second, the observational scale, which determines the degree of environmental heterogeneity associated with that scale. At the largest possible scale, that of the entire planet, all species on Earth co-occur, despite all kinds of environmental differences. As we move to finer spatial scales, and segment space into a definite number of sites, common species will tend to co-occur with many others simply because they are widespread across those sites. On the contrary, rare species, which may appear in just fewer sites, will tend to co-occur with far fewer others. A poor collection of local sites from a global sampling will probably include a high degree of environmental heterogeneity and may fail to integrate enough rare species, thus failing to reproduce more general patterns of species co-occurrences. It is essential, then, to work with enough local sites across rather uniform environmental conditions and standardize our measurements across them.

A key piece of information, therefore, is how frequently a species occurs across all local sites—its total occupancy—as well as how large an area these sites collectively cover, since this determines environmental heterogeneity and thus, at least in an operational sense, defines the species pool. Moreover, if we extend the temporal window of observation, the probability that any two species co-occur will also typically increase. Hence, both temporal and spatial observational scales—together with the degree of commonness or rarity of species and their tolerance to environmental heterogeneity—can strongly bias the apparent tendency of species to be found in association with one another.

Our aim is to account for these influences and to develop a species-level index that minimizes such biases. Specifically, building on our co-occurrence–occupancy curve, we define the Species Association Index (SAI), an occupancy-standardized measure of the tendency of a species to associate with others. This index allows comparison of the tendency of two species to associate with others, even when their total occupancies differ greatly. Our index performs best when environmental heterogeneity among sites is low, because it relies on a null hypothesis that assumes sites are equivalent.

In this study, we begin by presenting the empirical co-occurrence–occupancy curve (the *M*-plot) for a simple hypothetical community. We then show the theoretically expected curve under our null model, which assumes site equivalence and independence among species. We define the strength of species association as the number of species with which a focal species co-occurs in at least one site across the entire network. This quantity is then standardized through the introduction of the Species Association Index. Next, we examine two empirical case studies and construct two *M*-plots using data from Mediterranean mediolittoral habitats (MMH) and tropical rainforest trees on Barro Colorado Island (BCI), from which we also derive a Species Association Index (SAI). Both *M*-plots display a similar characteristic shape. Furthermore, we observe that the majority of species are consistent with the null model. We subsequently derive, in analytical form, the expected co-occurrence–occupancy relationship under both colonization–extinction dynamics and more general occupancy distributions. Finally, we summarize the main advantages of our approach and discuss its limitations.

## Methods

### The empirical co-occurrence-occupancy curve

First, we introduce the *M*-curve (the *M* from e*M*pirical). This is an empirical curve, a community-level pattern based on community occupancy data, which can be plotted from species occurrences across a number of sites distributed in a given area. Usually, occupancy data are presented in a table:

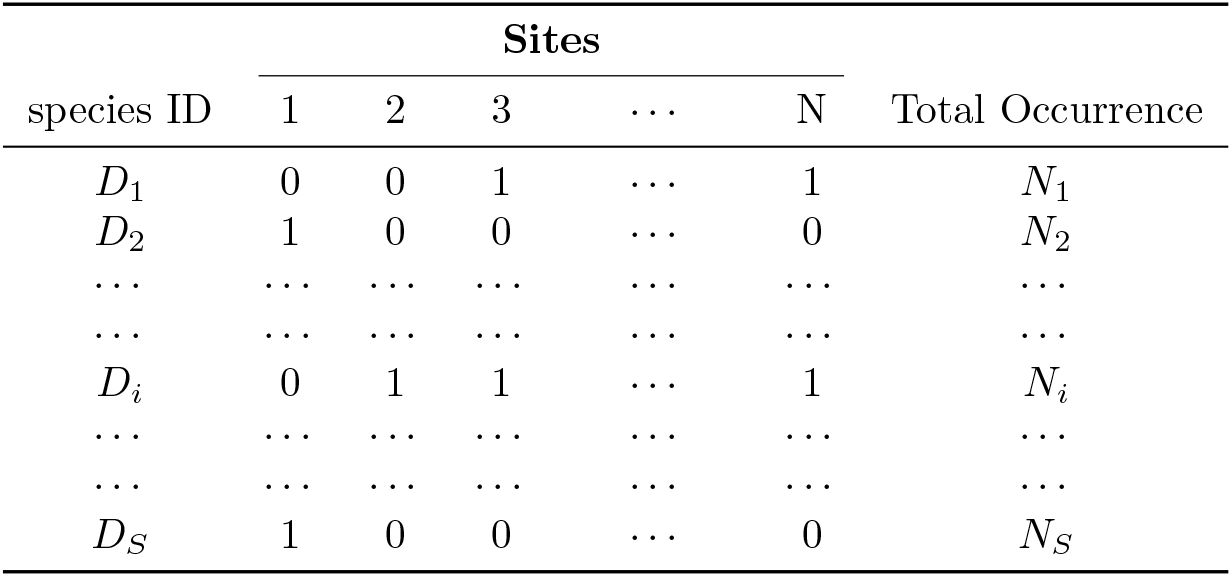

where a number of *S* total species are found across *N* total sites. Notice that we have used lower case *n*_*i*_ to identify species *i* and capital *N*_*i*_ to count the total occurrence, i.e., the total number of sites in which species *i* is present.

A simple way to quantifying species association is to count the number of instances in which a given species co-occurs with other species. Here, we define two species as *co-occurring* if they are found together in at least one site. As noted above, this measure is influenced by a species’ overall frequency of occurrence—its total occurrence (or occupancy). Accordingly, this association metric can be examined as a function of a measure of species occurrence, such as its global occupancy.

Based on this initial definition, and as a measure of the propensity of a given species to appear associated with others, we further define its association strength as the number of distinct species with which a focal species *co-occurs* across all sampled sites. When this quantity is evaluated for every species in a community, it yields what we term the empirical co-occurrence–occupancy curve, or the *M* plot (see Fig. 1). For simplicity, when plotting each species’ association strength against its global occupancy, we typically label the *y*-axis simply as “co-occurrences.” As discussed, the more broadly a species is distributed across sites—that is, the greater its occupancy—the greater its association strength will tend to be. It is important to note that the association strength of a species is only defined in the context of a network of local sites.

**Figure 1.**
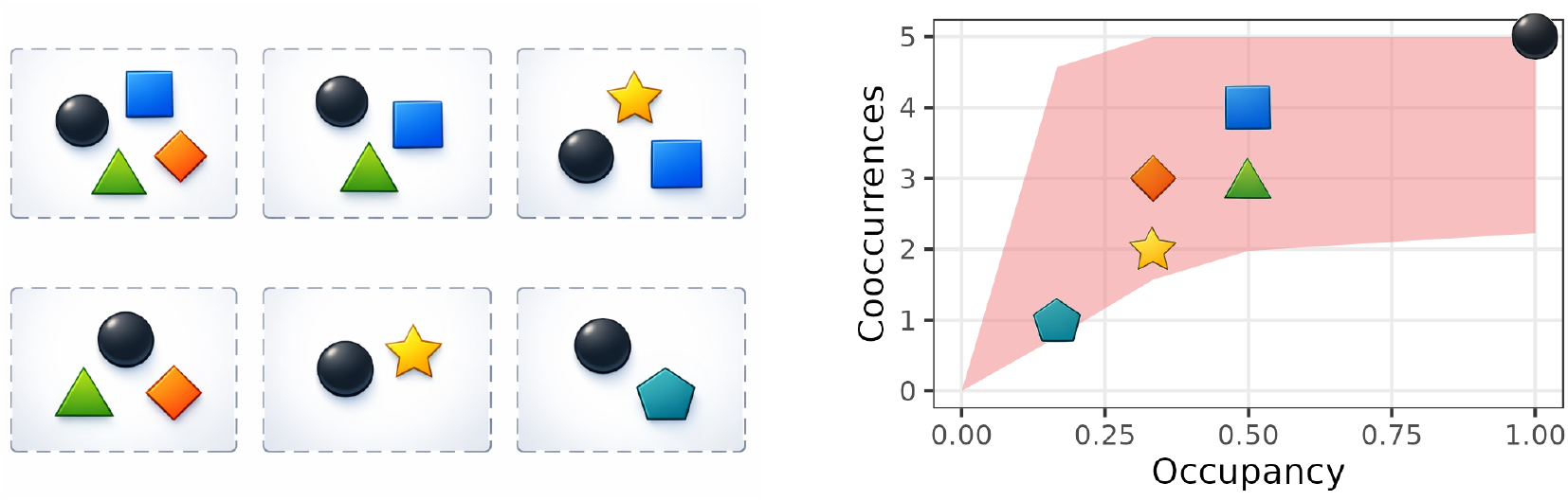
The M-curve. *Left*, six imaginary communities with a total of six species (symbols). We are interested in how many species a focal species co-occurs with in the metacommunity. The circle species co-occurs with every other species, whereas the pentagon species only co-occurs with the circle. *Right*, the empirical curve (*M*-curve) of the metacommunity. Note that the triangle species, even though it has the same occupancy as the square species, has fewer co-occurrences. The M-curve can be predicted based on an empirical species-by-site matrix. We provide means to calculate the expected value and 95% distribution of the co-occurrences (red ribbon) in this contribution.

Since species occupancies can be scaled between 0 and 1, in order to be able to compare across systems, we can also re-scale the *y* axis. To be more precise, on the *x* axis, we plot species occupancies, this is, total occurrence of a species across sites divided by the number of sites:

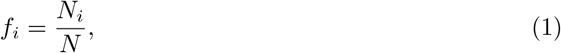

as you can see in Fig 1, and, on the *y* axis, we may plot the number of the observed association or co-occurrence level a of a given species, *A*_*i*_ (as in Fig 1a), divided by the maximum number of potential co-occurrences, *S*−1. We call this quantity species relative association (or co-occurrence) strength of the *i*-th species.

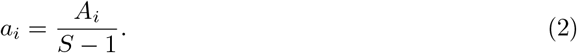

## Results

### The Null Model

Let us assume now that species *i* occurs on *N*_*i*_ sites. Given that for the rest of species each of them occur certain number of times across all sites, what is the probability for the *i*-th focal species to co-occur with exactly *A* species among the rest of *S* − 1 potentially co-occurring species? Mathematically, we are interested in the distribution of the *A* random variable. In nature, this distribution will depend on a range of different factors affecting how the system of *S* species across the *N* sites has ben assembled. However, we assume minimal hypotheses in the absence of mechanisms and species idiosyncrasies of any kind. Our *null* hypothesis is based on two assumptions:

1. **Species independence**. Species are placed across sites without feeling the influence of each other.
2. **Random placement (or Site equivalence)**. Species do not have any preference for particular sites, this is, there are not sites that tend to attract more species than others. Sites are equivalent.

Interestingly, in 2013, Veech gives the probability that two species can co-occur at exactly *n* number of sampling sites given that each species occurs at *N*_1_ and *N*_2_ number of sites out of a total of *N* under, precisely, the *null* hypothesis detailed above. Under random and independent placement, this probability is:

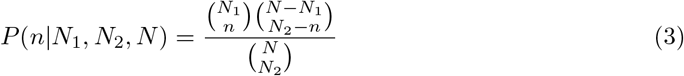

It can be shown that this equation is symmetric under swapping *N*_1_ and *N*_2_ and equivalent to that provided by Veech [35].

It follows that the probability of species 1 and 2 co-occurring in at least one site is given by:

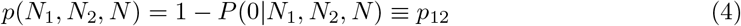

Therefore, in a system defined by *S* species independently and randomly placed on *N*_1_ to *N*_*S*_ total sites out of a total of *N*, we can express the average number of co-occurrences of a focal species *i* with the rest *S* − 1 species as:

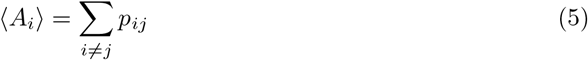

where, according to Eq (4) and (3), the *p*_*ij*_s can be written as:

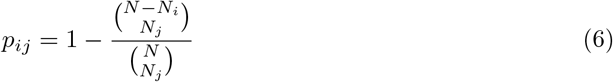

Therefore, the average version of Eq. (2), this is, the expected value of the relative association (or co-occurrence) level of a focal species *i* under the two null model assumptions stated above, becomes:

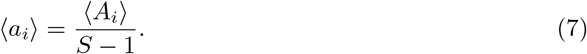

Notice that the quantity ⟨*A*_*i*_ ⟩ represents the expected value of a random variable *A*_*i*_ that follows the species co-occurrence distribution when the species *i* is present in *N*_*i*_ sites, and the rest of them (*j* ≠ *i*) in *N*_*j*_ sites. As we plot this metric for every species in a community with total occurrences, *N*_1_, *…, N*_*S*_ across a total of *N* sites, we have the expected co-occurrence-occupancy curve under the *null* hypothesis (see thick black line in Fig 2). We ask now about the variance of *A*_*i*_, this is, how variable the number of co-occurrences a given focal species can be. It turns out that the variance of this distribution can be also calculated analytically in terms of the average given before and the probabilities in Eq (6) (see appendix A.1):

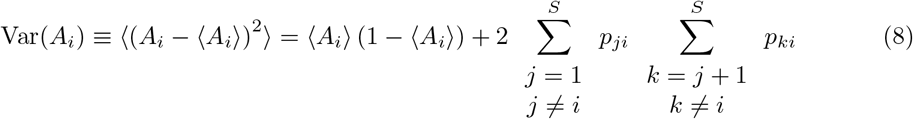

**Figure 2.**
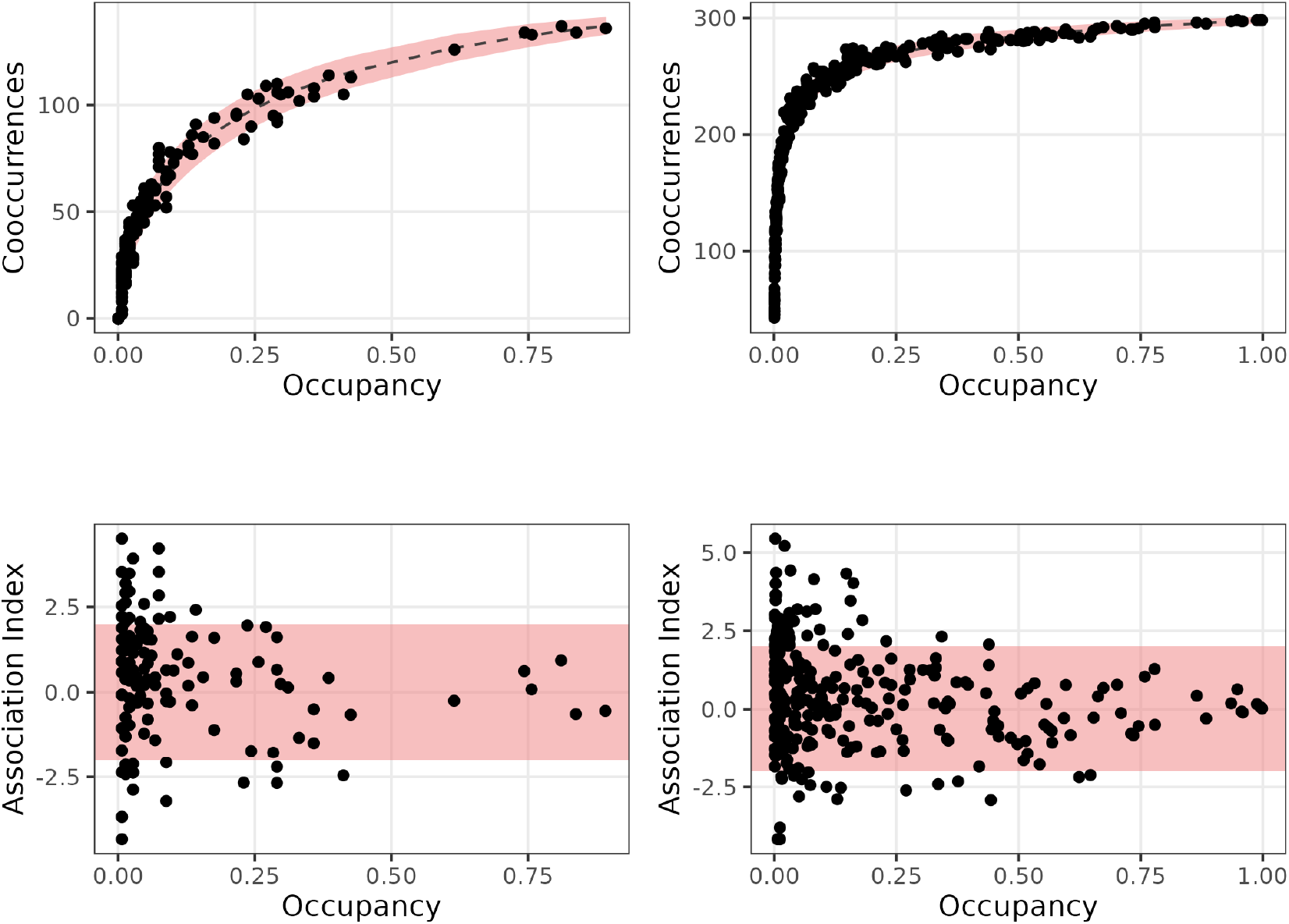
The co-occurrence-occupancy plot. Here we compare two distinct communities. On the the two panels on the left, we plot the *M* curves for the MMH, and on the panels on the right, we plot the same curves for the BCI rain forest trees. In the upper plots (a and b), each point represents a given species, where the *x* coordinate represents its global occupancy across sites, and, the *y* coordinate is the number of species that species happens to co-occur with, its co-occurrence or association level The red ribbon indicates the 95% confidence interval of the expected cooccurences, while the black dashed line indicates the expected cooccurrence. In the lower plots (c and d), the *x* axis is the same, but the *y* axis represents the association index for every species in the system. Points outside of the red zone are considered to deviate significantly from expectation.

### The Species Association Index (SAI)

Because the total number of occurrences of species across sites influences their association strength, we apply a *z*-score standardization to make association strengths comparable among species that vary in their overall occurrence (or occupancy). This motivates the definition of the Species Associativity Index (SAI), *s*_*i*_, for species *i* as:

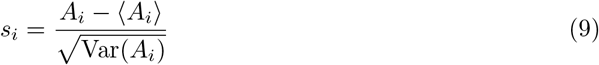

where *A*_*i*_ is the number of observed co-occurrences of species *i* with all other species, out of the *S* − 1 species with which it could potentially co-occur. As we have shown, this index can be calculated analytically. In theory, its values may range from −∞ to +∞, although in practice it typically lies between -5 and +5 in most communities. When a species co-occurs with others far more frequently (high association index) or far less frequently (low association index) than expected under the *null* hypothesis, this suggests that non-random processes are driving the corresponding positive or negative associations. In addition to the empirical curve (Fig. 2, top panels), we can also plot the association index for each species in the system (see Fig. 2, bottom panels).

### The theoretical co-occurrence-occupancy curve under generic occupancy distributions

Thus far, we have derived the expected co-occurrence–occupancy relationship for a set of *S* species with specified total numbers of occurrences, *N*_1_, *…, N*_*S*_, across *N* sites, under a null model based on random placement and species independence. This relationship represents the average curve that would be obtained by repeatedly generating species occurrences at random across the *N* sites. In each randomized replicate, every species is constrained to occur exactly *N*_*i*_ times, but these occurrences are allocated to a different randomly selected subset of sites drawn from the full set. This randomization procedure enables the estimation of percentile values for the distribution of the species association level, *A*_*i*_, corresponding to each empirical total occurrence (see Fig. 2), under the assumed null hypothesis.

As previously discussed, the distribution of the association strength, *A*_*i*_, for a given species depends jointly on its own total occurrence and on the total occurrences of the remaining *S* − 1 species across the *N* sites. More generally, the manner in which occupancy is distributed among species strongly influences the shape of the co-occurrence–occupancy relationship.

To illustrate this, we first generated two markedly different occupancy distributions (top row of Fig. 3). We then randomly sampled *S* species occupancies, *p*_*i*_, from each of these distributions, which defines *N*_*i*_ as the closest integer number to *p*_*i*_ *N*. Conditional on species *i* having a global occurrence *N*_*i*_, we randomly selected *N*_*i*_ sites from the full set of *N* sites. Repeating this procedure for every species allowed us to construct the co-occurrence–occupancy curve corresponding either to a single realization of this generative process or to its expected value over many realizations (bottom row of Fig. 3).

**Figure 3.**
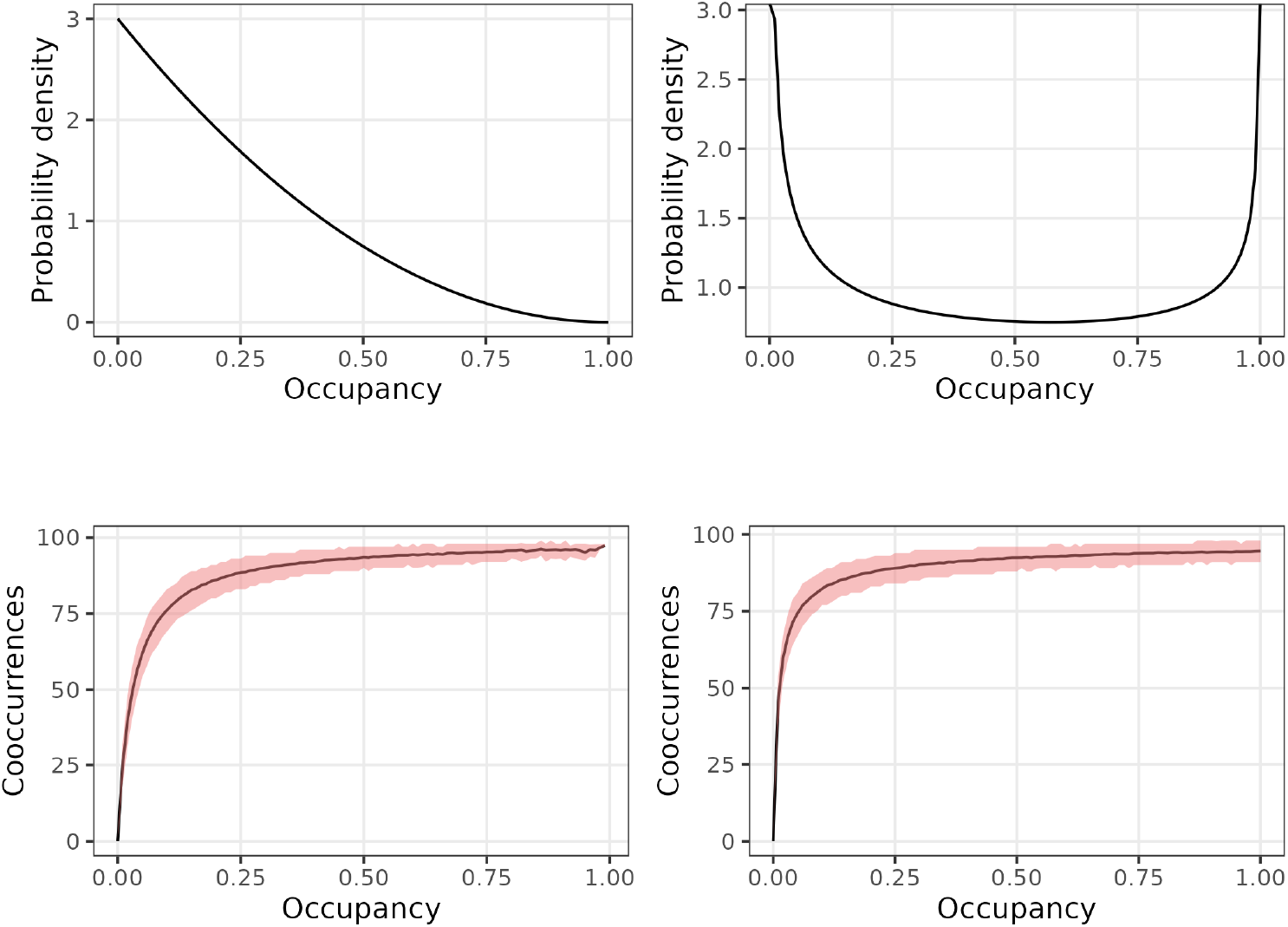
The occupancy distribution and the co-occurrence-occupancy relationship. Plots are generated from two beta functions with different *α* and *β* parameters. The upper plots are the probability density functions of the beta distribution for alpha = 1; beta = 3 (left), and alpha = 0.6, beta = 0.7 (right). From each of these two parameter sets, we obtained the occupancies for 100 species. Those were translated into presence/absence for a community with 100 sites. We repeated this procedure a 1000 times for each parameter set. The lower graphs represents the 90% distribution (red ribbon) and mean (black line) of all species realized occupancies vs. their co-occurrences.

**Figure 4.**
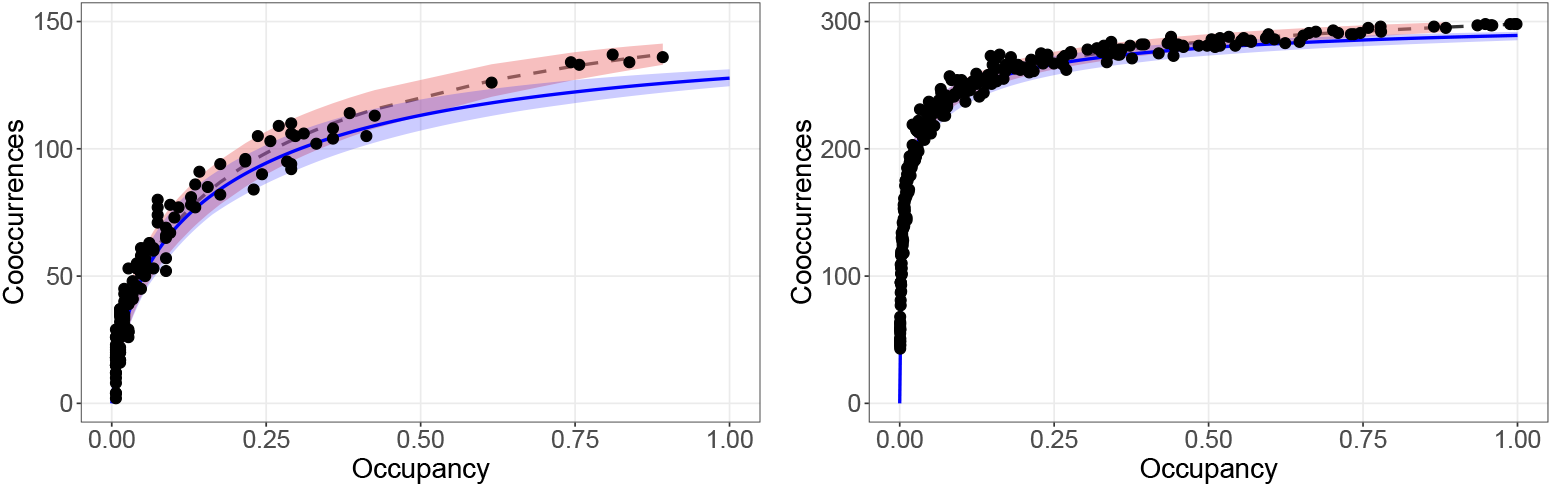
The co-occurrence-occupancy relationship of the case studies. Plots are generated from the treated data of the case studies. They correspond to the comparison of the empirical data (black dots) with the comparison of the null model (grey dashed curve), and the direct computation of Eq. (15) (blue curve) from the empirical data for: the Mediterranean rocky shore community (left) and BCI (right). The coral and blue ribbons correspond to the 95% confidence interval for the null model and for the computation of Eq. (15) from bootstrapping over 200 realizations, respectively.

It is important to note that this expected curve is more general than the one obtained previously, because the expectation is now taken with respect to a generic occupancy distribution, rather than over random permutations of a fixed vector of species total occurrences, (*N*_1_, *…, N*_*S*_). Notice also that Fig 3 has been generated under the additional assumption of species equivalence, since all species are drawn from the same occupancy distribution.

Now we show, more generally, that within our initial hypothesis, this is, under species independence and random placement, we can calculate the expected co-occurrence-occupancy curve under different hypothesis about the processes determining the species occupancy distribution. The first example assumes colonization-extinction dynamics and the second a typical general occupancy distribution.

#### Colonization-Extinction Dynamics

Under the assumption of species independence, a convenient approach to characterizing the occupancy of a species across a set of equivalent sites is to model its dynamics using a colonization–extinction framework [2]. Within this framework, colonization (*c*) and extinction rates —and hence equilibrium (stationary) occupancies—can be inferred from temporal presence–absence data [26]. Accordingly, the expected occupancy of species *i* is given by:

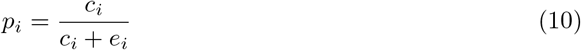

In this framework, we characterize each species by its corresponding expected occupancy probability, denoted *p*_1_, *…, p*_*S*_, taking values in the interval [0, 1]. It is important to emphasize that this specification does not impose species equivalence. Under the assumption of species independence, the total numbers of occurrences across sites consequently follow independent, but distinct binomial distributions:

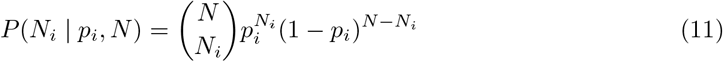

Recall that the probability that species *i* and *j* co-occur in at least one site is a function of their total numbers of occurrences, *N*_*i*_ and *N*_*j*_, across the complete set of *N* sites, see Eq. (4). In the present framework, *N*_*i*_ and *N*_*j*_ are random variables distributed according to the independent binomial laws specified above. Consequently, the expected association strength of species *i*, ⟨*A*_*i*_⟩, must be calculated by taking the expectation of *A*_*i*_ with respect to these binomial distributions, *P* (*N*_*i*_ | *p*_*i*_, *N*), accordingly.

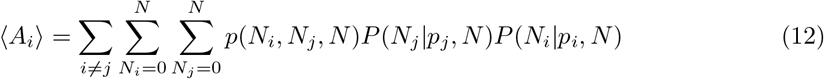

Note that the co-occurrence strength of a species remains a function of the total number of sites in the system, *N*, as well as the stationary expected occupancies of all species, *p*_1_, *…, p*_*S*_. These occupancies, in turn, are determined by the species-specific colonization and extinction rates.

#### A general occupancy distribution

In nature, uneven occupancy distributions are the rule [19]—most species are very rare and occur at a few sites, while few others are almost ubiquitous—. A quite general distribution, with a support between 0 and 1, able to present this behavior is the the Beta distribution (see Fig 3, panel A). If we make the extra assumption of species equivalence, this is, we consider all species occupancies identically and independently distributed following the same occupancy distribution, then the expected association strength of a given species should consider the expectation weighted by the *S* independent occupancy distributions:

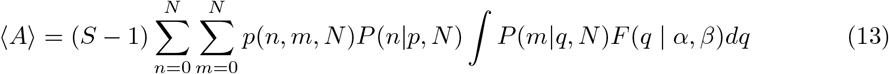

This expression is valid for a generic occupancy distribution *F* (*p*) as long as its support is between 0 and 1. This average value is a function of the occupancy, *p*, the parameters of the distribution, say *α* and *β*, and the total number of species, *S*, and sites, *N*. As a function of *p*, it corresponds to the theoretical co-occurrence-occupancy relationship under a generic (two-parametric) occupancy distribution, *F* (*p* | *α, β*). Interestingly, if this is the Beta distribution, which we argue it is a reasonable way to model species occupancies across sites (see Fig S2 and S3 in Supporting Information), it turns out that the expected curve can be expressed as a function of the occupancy, *p*, the total number of species, *S*, and sites, *N*, and the probability density parameters, *α* and *β*, in a quite condensed manner:

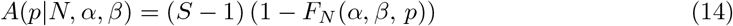

where the function *F*_*N*_ (*α, β, p*) is a well-known mathematical special function (see appendix A.2 for details). For a general distribution for the occupancies with density *F* (*q*)—not necessarily a Beta distribution—one can get a general expression, as follows,

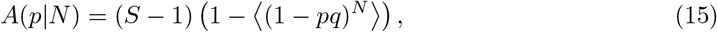

where 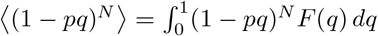. Importantly, this expression can also be evaluated, directly from the empirical occupancy distribution, as a sum over the observed occupancies, {*q*_1_, *…, q*_*S*_}. That is, for every value of *p*, we can approximate Eq. (15) by

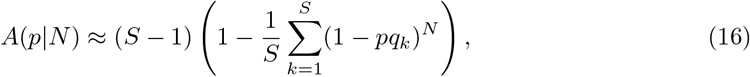

avoiding integration and any further assumptions on the theoretical occupancy distribution species should follow.

In Fig. S3 we can see the comparison of the empirical data and the results from the null model and the Eq. (15). The 95% confidence interval for the estimated curve for Eq. (15) is computed using bootstrapping [39]. Eq. (15) might lead to better than Eq. (14) estimates since its computation is independent of the assumption of the empirical distribution. A discussion on that for the empirical data studied in this work can be observed in Appendix A.3.

### Case studies

#### A rocky shore community

The Mediterranean mediolittoral habitats represent a subset of the main littoral communities from the rocky shores of Catalonia, concretely those thriving in a tiny fringe just at the sea level along a stretch of coastline of about 1100 Km. These communities were sampled between 2010 and 2012 along 143 hard-bottom rocky shores and man-made structures such as breakwaters and jetties. All visible flora and fauna found within the transects were recorded using a Braun-Blanquet [4] cover-abundance scale. The methodology has been described in different works (e.g. [6, 9]).

Co-occurrence and occupancy patterns in the MMH mostly followed the null model. Nevertheless, some species scored lower or higher in their association index (SAI) for very similar occupancy levels. Species from very simplified habitats, like those close to sandy shores or those creating crusts not colonizable by other species co-occurred very poorly, thus had a lower SAI than species with relative mobility (e.g. some gastropods), the epibionts, or well-known opportunists. Interestingly, the greatest departures from null expectations were found for species characteristic of other levels of the littoral fringe, this is, species that may not belong to the same mediolittoral habitat. This may help defining accurately the limits and subunits or a natural community.

#### A rain forest community

The BCI 50ha forest dynamics plot was specifically designed to observe community-level processes shaping tree richness and to understand the long-term dynamics of rare and abundant species across spatial scales and life-history stages [17]. All trees with a minimum diameter at breast height of 1 cm has been surveyed every five years since 1982, following standardized protocols. We used the 2015 survey [7], considering as sites its 1250 quadrats (20×20m), which comprised 299 different tree species.

Here, we found that the co-occurrence and occupancy patterns mostly followed our proposed null model, which is not a surprise given that this system is an inspiration for neutral theory [16]. Interestingly, the association index has some variability. One potential cause is that the tree community may be structured by two demographic trade-offs, a growth-survival one, and a stature-recruitment trade-off [29, 30]. Actually, the association index depended on these two axes (Fig. S1), the stature-recruitment one modulated by wood density (p-value *<* 1e-07, *R*^2^ = .12). This result further supports that both demographic axes control forest community structure and dynamics.

## Discussion

Species co-occurrence remains a central construct in community ecology, not as a crude proxy for interaction but as a statistically robust and conceptually versatile framework for investigating community assembly, structural organization, and emergent properties. Initial tendencies to interpret spatial association as direct evidence of biotic interaction have been tempered by recent methodological and conceptual critiques [3, 22]. These contributions have delineated the inferential limits of co-occurrence data and enhanced analytical rigor, while leaving the core relevance of co-occurrence approaches intact.

Contemporary research demonstrates that co-occurrence patterns can uncover higher-order structural attributes of ecological systems, including network scaling relationships [12] and non-neutral community modules [21]. Methodological developments have substantially expanded the scope of co-occurrence analyses, ranging from machine-learning–inspired embedding techniques [32] to the incorporation of environmental DNA data into co-occurrence network frameworks [33]. In parallel, empirical and conceptual re-evaluations of the definition of ecological “communities” [31, 36], together with syntheses on emerging community data streams [15], underscore the role of co-occurrence as a foundational bridge between empirical observations and ecological theory. Rather than being rendered obsolete, co-occurrence analysis has evolved into a more nuanced, statistically grounded paradigm that continues to inform debates on ecological neutrality, niche differentiation, and the overall architecture of ecological communities.

In this context, our paper spotlight a simple yet largely overlooked community-level pattern: the co-occurrence–occupancy curve. This curve captures how strongly a species tends to show up with others—co-occurring at the same sites—relative to its overall occupancy across all sites. We rigorously define this empirical relationship—the *M* curve—and derive its expected shape under a random null model that assumes equivalent sites and independent species. Building on this foundation, we introduce the Species Association Index (SAI), a new occupancy-standardized metric that quantifies how prone a species is to appear in association with others, independently of how frequent it is overall. By decoupling association from raw occupancy, the SAI allows meaningful comparisons among species with very different commonness and supplies a clear, quantitative benchmark for detecting and interpreting departures from neutral expectations.

The simplest null model we introduced assumes species independence and random placement. Under this null hypothesis, species see the sites as environmentally homogeneous. This does not mean that sites are just identical replicates of each other, but rather that their inherent differences (due their location and distinct environmental characteristics) do not influence the establishment of the species. Therefore, tolerant, generalist species will follow this model more closely that strongly specialized ones.

We have also extended our results when species are not simply randomly placed on the sites, but follow certain independent dynamics. For instance, the predicted co-occurrence-occupancy relationship (and the expected SAI for each species) can also be linked to a useful stochastic colonization-extinction framework [2, 26, 27, 18], and to any dynamic community model predicting the species occupancy distribution [25, 19].

When applying our approach to data, it is important to recall we have defined a metacommunity (a set of communities on a network of sites) as sharing similar abiotic conditions, resource availability, and trophic structure within a given spatial domain. Following Vellend [37], we have considered a *horizontal* community, this is, a group of organisms co-occurring in a particular place and time and sharing a broadly similar ecology. To stick to this constraint, we have avoided gradients of resource availability and other abiotic conditions, and focused on primary producers and sessile or weakly mobile organisms, rather than on species that readily move along different environmental gradients. Among other reasons (see below), we have found that departures from null expectations may arise from the existence of high heterogeneity across sites or even inadequate sampling designs that combine ecologically incomparable systems, thereby undermining meaningful inference.

We have illustrated our approach using two contrasting systems—tropical rain forest trees on Barro Colorado Island and organisms from Mediterranean rocky shores. Through our analysis, we have identified the ecological mechanisms underlying deviation from neutrality in these systems. Explanations range from simple distributional effects to more complex species- and community-level processes. We have quantified the ability of a species to co-occur with others through its species association index. A species has a high score when it co-occurs significantly more often than expected on the basis of its occupancy, and a low score when it co-occurs less than expected. This formulation has allowed us to order species along a continuous association axis, from highly *sociable* to comparatively *solitary* taxa. We have discussed the ecological implications of this ordering at both species and community levels. In particular, we have proposed that species with higher tendency to appear associated with others are both structural or engineering species, and epibionts, relatively mobile across life-history stages, opportunistic in resource use, and characteristic of more complex communities. Conversely, species that generate or occupy structurally simplified or environmentally impoverished patches may score low on the association axis.

By reframing co-occurrence through the lens of occupancy, we provide a simple, transparent baseline for asking a fundamental ecological question: who tends to live with whom, and why? The co-occurrence–occupancy curve and the Species Association Index (SAI) together disentangle frequency from association, allowing neutrality to serve not as an assumption but as a testable benchmark. In doing so, this framework opens a tractable path toward detecting the signatures of interaction, environmental filtering, and historical contingency in the architecture of ecological communities.

## Author contributions

Vicente J. Ontiveros contributed the figures, curated the data, and developed the *R* code and package used to evaluate the curves, species association indices, and plots. Simone Mariani collected samples from the mediolittoral habitats in Catalonia and brought the concept of the *M*-plot to the authors’ attention. Alberto Megías participated in the mathematical formulation of the models and in obtaining the estimates for the theoretical curves and plots, while Leonardo Aguirre and José A. Capitán verified and enhanced the internal consistency of the article’s mathematical derivations. David Alonso drafted the initial version of the manuscript and the supporting information and consistently inspired perseverance and enthusiasm within our group. All authors contributed equally to the intellectual development of the work and to the editing of the manuscript.

## Acknowledgments

VJO was funded through grant PRIORITY (PID2021-127202NB-C22) awarded to JAC. LA was funded thorugh grant UNIQUE (PID2021-127202NB-C21) awarded to DA. Both grants were funded by MCIN/AEI/10.13039/501100011033 and “ERDF. A way of making Europe”. We gratefully acknowledge the many contributors responsible for the long-term sampling and careful data curation of the 50-ha plot on Barro Colorado Island. The basics of this research date back to the MSc thesis of Arnau Luke Dedeu Dunton with whom we shared fruitful thoughts about the mediolittoral communities from Catalonia and species co-occurrences. We also dedicate this work to E. Ballesteros, whose support, inspiration, and deep love for the benthic communities of the Mediterranean rocky shores have been a constant source of motivation.

## A Supporting Information

### A.1 The variance under the *null* hypothesis

Since the *null* hypothesis assumes a process of independent and random placement of *S* species on exactly *N*_1_ to *N*_*S*_ total sites across a total of *N* sites, the quantity *A*_*i*_ can be regarded as a random variable counting the number of co-occurrences in at least one site of focal species *i* with the rest of *S* − 1 potentially co-occurring species. Due to species independence, this variable can be written as a sum of indicator random variables:

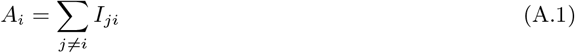

where *I*_*ji*_ is a boolean random variable, this is, it takes the valule of 1 when species *i* and *j* co-occur in at least one site, and 0 otherwise:

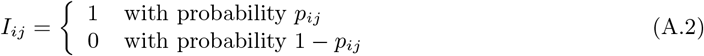

where the probabilities *p*_*ij*_ are defined according to Eq (6) in the main text. Since we are interested in the variance of *A*_*i*_, and this random variable has been carefully defined in terms of a sum of random variables whose distributions are also defined (see Eq. (A.2) above), we can proceed to calculate the variance in the usual way:

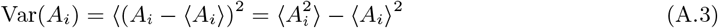

First, we calculate the expected value of *A*_*i*_. Again species independence simplifies the calculation very much. So we recover Eq (13) in the main text:

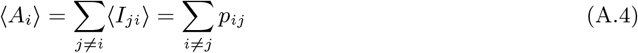

Then, we calculate the expected value of 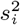.

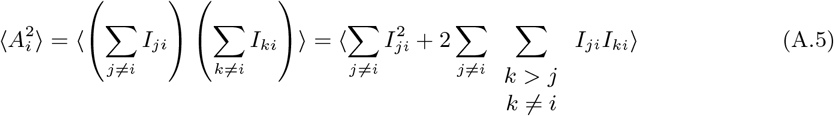

Since the expected value of a sum of random variables is always the sum of expected values, we can calculate the two terms within brackets above separately. First:

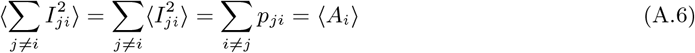

and second:

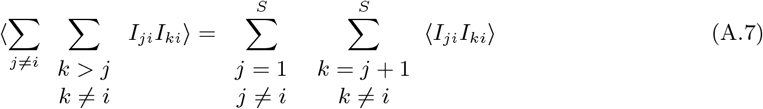

Finally, the expected value of the product of two boolean random variables is given by:

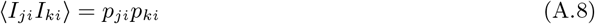

Therefore, if we introduce all this in Eq (A.3) for the variance of *s*_*i*_, we finally obtain the expression given in the main text (see Eq. (8)), as we wanted to show:

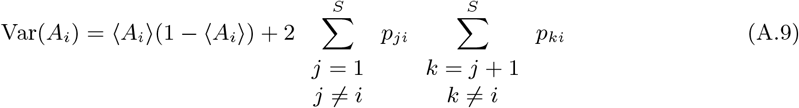

### A.2 The species occupancy distribution and expected co-occurrence-occupancy relationship

Here, we derive Eqs. (14) and (15) of the main text. Let us start from Eq. (13),

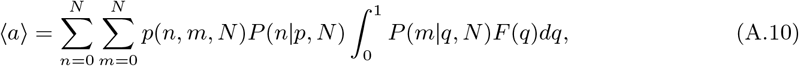

where *a* = *A/*(*S* − 1) and we are not going to assume an explicit form of the distribution *F*. Using the expressions for *p*(*n, m, N*) we have that,

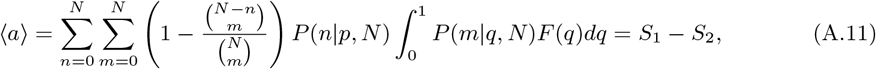

with,

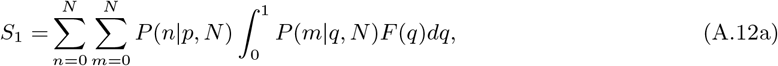

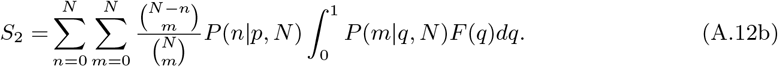

From normalization of the binomial distribution 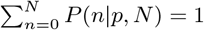, then

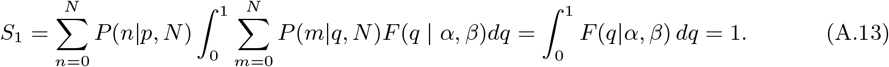

Note that the former result is independent of the distribution *F*. Here, we just use the fact that the support is in the [0, 1] interval. Hence, ⟨*a*⟩ = 1 − *S*_2_. Moreover, we can express

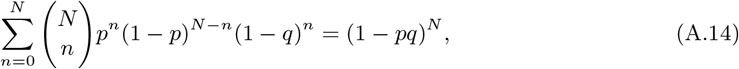

by using the Newton’s binomial rule. Therefore,

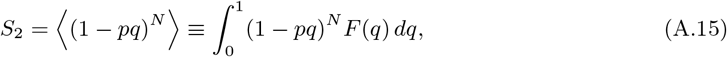

which is general for every distribution and implies

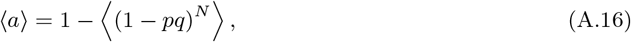

resulting in Eq. (15). This expression is quite useful; since it can be expressed as an average, it can be computed as a sum over the observed distribution of occupancies. That is for every value of *p*, given the empirical set of occupancies {*q*_1_, *…, q*_*S*_ } we can approximate

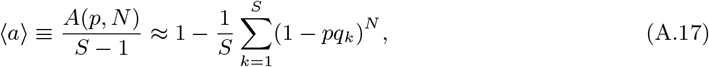

avoiding any integration and directly from the empirical dataset.

In the particular case where *F* (*p*) = *F* (*p* | *α, β*) stands for the Beta distribution,

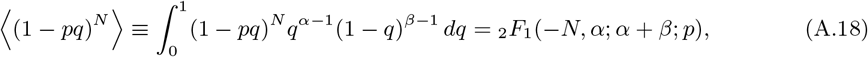

and, then,

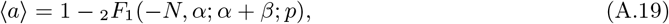

which implies the form of Eq. (14).

#### A.2.1 Expected value of association index

The expected value of association index, where *p* is drawn from the same Beta distribution *F* as the other species’ *q*, can be expressed through (polynomial) hypergeometric functions in a similar way, namely

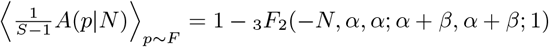

This can be computed as the *N*-th iteration *a*_*N*_ of the recursion

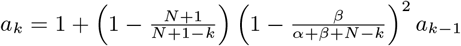

with *a*_0_ = 1.

### A.3 Parameter estimation for Beta distribution

To fit the empirical points for the co-occurrence-occupancy curves to Eq. (14) assuming a Beta distribution for the occupancies we elaborated two strategies. First, we computed the maximum-likelihood estimates (MLE) for the parameters of the Beta distribution using the empirical data for occupancies. Second, we performed a nonlinear fit from the `curve_fit` function of the `optimization` package of SciPy [38]. The maximum-likelihood estimation of the first strategy is elaborated in more detail in the next section.

#### A.3.1 MLE of the Beta distribution

The probability density function for a Beta-distributed random variable *x* ∈ [0, 1] is defined as follows

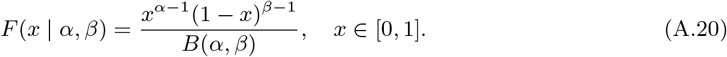

For a sample of *n* i.i.d. observations 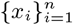—here, the occupancies,—the log-likelihood function 𝓁is defined as

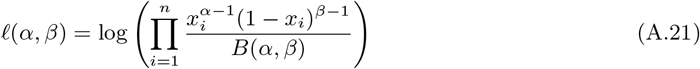

which after some algebra reads

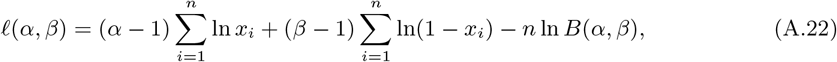

where *B*(*α, β*):= Γ(*α*)Γ(*β*)*/*Γ(*α* + *β*) is the Beta function and Γ the Gamma function. The MLE are based on the maxima of the log-likelihood function, hence we need to compute the partial derivatives of it, which read

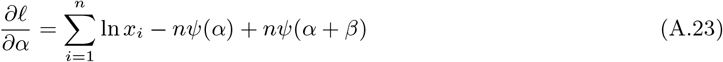

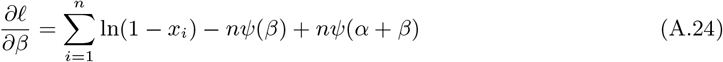

where 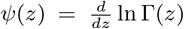 is the digamma function. The MLE pair (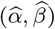) fulfill that 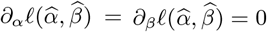 and the condition of maxima for a bi-dimensional function. These estimates cannot be expressed in a closed form in terms of known functions and solutions for this problem are obtained from numerical methodologies. Here, they are numerically estimated using the `stats` package of SciPy [38].

#### A.3.2 Comparison of the different estimates

In Figs. S3 we show the comparison of the two proposed methods for estimating Eq. (14) from the empirical datasets with respect to the estimate of Eq. (15). We can observe that the two estimate methods for Eq. (14) almost coincide in their confidence interval, but deviates more to the data than Eq. (15) does.

A possibility of the deviation might be that the Beta distribution might not be the best fit to the data. In Fig. S2, we show the MLE fits of the empirical histogram for the Beta distribution. The larger deviations from the MLE probability density to the empirical histogram are located in the tails of the distribution close to the occupancy 1 as well as occur for the estimates of the co-occurrence-occupancies estimated curves in Fig. S3. In Table 1 there is a summary of the 95% confidence intervals for the parameters of the Gamma distributions computed from bootstrapping [39].

**Table 1.** 95% confidence interval for the parameters of the Beta distribution for the empirical data from MLE of the empirical distribution and the nonlinear fit of Eq. (14).

|  | MLE | Nonlinear fit |
| --- | --- | --- |
| MMH | $CI_{95\%}(\alpha) = (0.445, 0.635)$ | $CI_{95\%}(\alpha) = (0.440, 0.483)$ |
| | $CI_{95\%}(\beta) = (2.402, 6.169)$ | $CI_{95\%}(\beta) = (3.406, 3.770)$ |
| BCI | $CI_{95\%}(\alpha) = (0.338, 0.431)$ | $CI_{95\%}(\alpha) = (0.397, 0.401)$ |
| | $CI_{95\%}(\beta) = (1.15, 1.98)$ | $CI_{95\%}(\beta) = (1.70, 1.80)$ |

## Supplementary Figures

**Figure S1.**
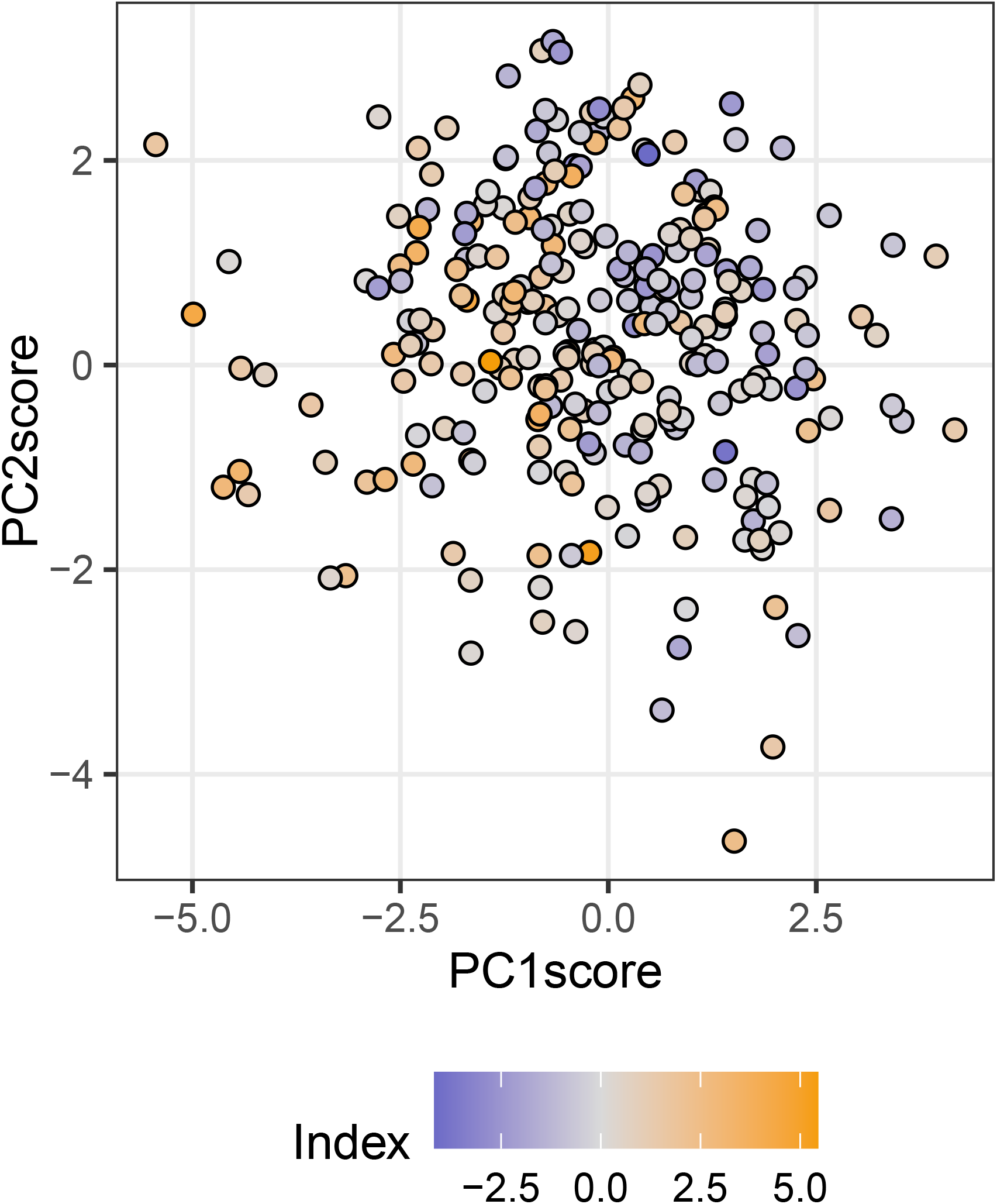
The Association Index is influenced by demographic trade-offs. The x-axis corresponds to a growth-survival trade-off, while the y-axis does it to a stature-recruitment one. The position for each species was extracted from [30]. Although noise is present, we observe that slow, tall species tend to be less associated with other species than fast, small recruiters. We estimated that the association index had the following relationship with demography and wood density: *I*_*A*_ = −.315 * *PC*_1_ − .729 * *PC*_2_ + 1.032 * *PC*_2_ * *wd*.

**Figure S2.**
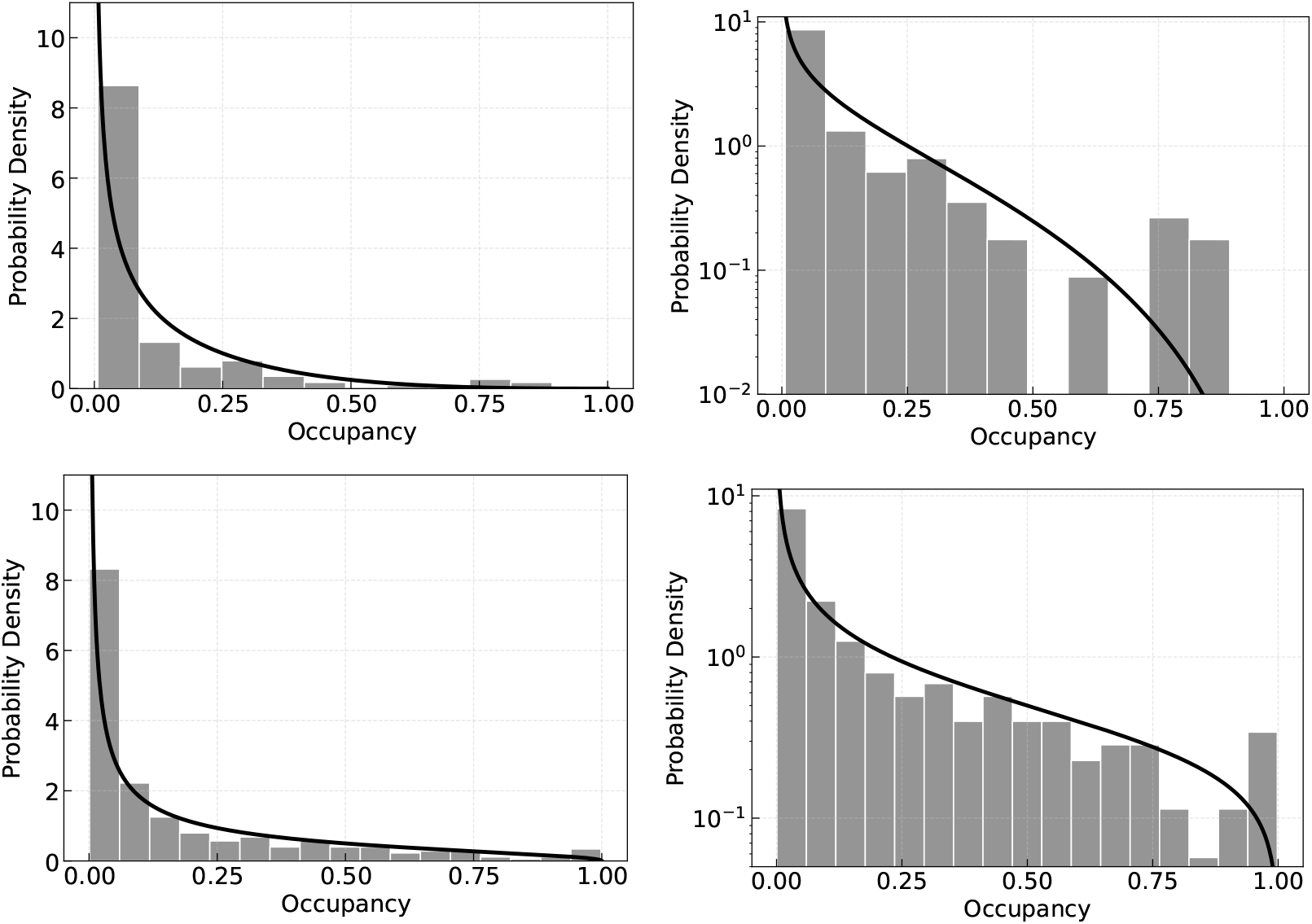
Probability densities from empirical data. Empirical probability density histograms (grey bars) and fitted Beta distribution density functions (black curves) for the MMH (top) and BCI (bottom) datasets. Results are shown on linear (left) and logarithmic (right) scales, with Beta parameters derived via MLE.

**Figure S3.**
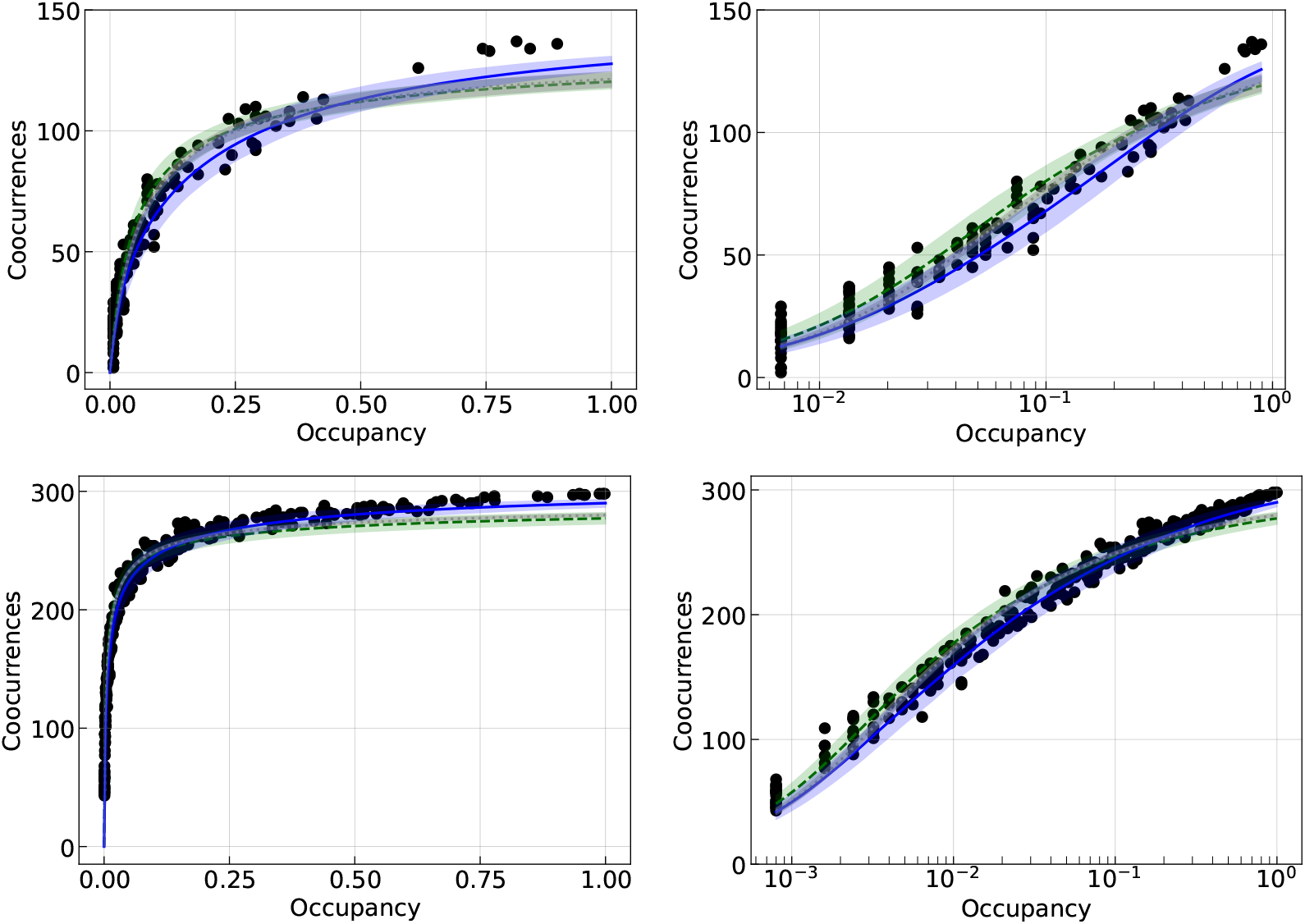
The co-occurrence-occupancy curve fits. Comparison of empirical data (black circles) with three theoretical models: (i) the expectation from Eq. (15) (blue curve and ribbon), (ii) Eq. (14) using Beta distribution parameters derived via MLE from empirical histograms (green dashed curve and ribbon), and (iii) a direct nonlinear fit of Eq. (14) (grey dotted curve and ribbon). All ribbons indicate 95% confidence intervals. Results are shown for MMH (top) and BCI (bottom) datasets in both linear (left) and logarithmic (right) scales.

